# Leveraging Allele-Specific Expression for Therapeutic Response Gene Discovery in Glioblastoma

**DOI:** 10.1101/2021.06.22.449493

**Authors:** Arko Sen, Briana C. Prager, Donglim Park, Zhe Zhu, Ryan C. Gimple, Jean A. Bernatchez, Sungjun Beck, Alex E. Clark, Jair L. Siqueira-Neto, Jeremy N. Rich, Graham McVicker

## Abstract

Glioblastoma is the most prevalent primary malignant brain tumor in adults and is characterized by poor prognosis and universal tumor recurrence. Effective glioblastoma treatments are lacking, in part due to somatic mutations and epigenetic reprogramming that alter gene expression and confer drug resistance. Here, we interrogated allele-specific expression (ASE) in 43 patient-derived glioblastoma stem cells (GSCs) to identify recurrently dysregulated genes in glioblastoma. We identified 118 genes with recurrent ASE preferentially found in GSCs compared to normal tissues. These genes were enriched for apoptotic regulators, including Schlafen Family Member 11 (*SLFN11*). Loss of *SLFN11* gene expression was associated with aberrant promoter methylation and conferred resistance to chemotherapy and poly ADP ribose polymerase inhibition. Conversely, low *SLFN11* expression rendered GSCs susceptible to the oncolytic flavivirus Zika, which suggests a potential alternative treatment strategy for chemotherapy resistant GBMs.

## INTRODUCTION

Glioblastoma ranks among the most lethal of human malignancies with current therapies only offering palliation (1). Reasons for treatment failure are myriad, with tumor heterogeneity at the genetic and transcriptional levels contributing to the malignancy of glioblastoma (2,3). Glioblastoma displays a functional cellular hierarchy with stem-like, self-renewing glioblastoma stem cells (GSCs) residing at the apex (4,5). GSCs contribute to resistance to chemotherapy and radiotherapy, neoangiogenesis, invasion into normal brain, and escape from the immune system (6-8). Therefore, targeting GSCs may improve current glioblastoma management and extend the lives of patients.

Although glioblastoma is one of the most deeply characterized solid tumors, precision medicine has not significantly benefited most neuro-oncology patients. Most studies and targeted therapeutic strategies have thus far focused on protein-coding mutations. However, many important mutations lie in non-coding DNA where they function by perturbing gene regulation. Non-coding mutations likely help drive glioblastoma tumorigenesis and drug resistance but are more challenging to identify and impact gene regulation in many different ways. Gene dysregulation can be caused by copy number alterations (CNAs) (9-11), as well as by mutations that affect splice sequences (12), untranslated regions (UTRs) (13), insulators (14,15), promoters (16,17), and enhancers (13,18). Moreover, genes are often regulated by multiple enhancers, which can be located hundreds of kilobases away from their targets (19). Regulatory mutations are, therefore, diverse, and frequently spread over very large regions of the genome. As a result, standard recurrence analyses that identify driver mutations in coding sequences are unlikely detect many important regulatory mutations in cancer genomes. Alternate approaches to discover regulatory mutations can by stymied by the myriad of non-coding mutations whose function is difficult to predict from sequence alone. Rather than relying on interpretation of non-coding mutations, unbiased identification of genes with altered regulation can pinpoint functionally important genes that are unlikely to be discovered through other methods.

Here, we leveraged allele-specific expression (ASE) as a new approach to interrogate recurrently dysregulated genes in glioblastoma. ASE measures the difference in expression between two alleles of a gene, by utilizing reads mapped to heterozygous sites (20,21). Unlike standard RNA-seq expression levels, ASE is particularly sensitive to cis-acting mutations because it is generally unaffected by trans-acting or environmental effects that impact both alleles equally. Thus, a major advantage of ASE is that it can identify genes that are dysregulated by cis-acting regulatory mutations, even when the specific identities of the regulatory mutations are unknown. However, ASE is not solely caused by somatic mutations and can also result from common germline polymorphisms (22,23), imprinting (24), or random monoallelic expression (25). Therefore, to discover genes that are dysregulated in cancer, the frequency of ASE in disease samples must be compared to a panel of normal samples. This approach has recently been used to identify new pathogenic genetic variants in muscle disease (21) and oncogenic mutations in T cell acute lymphoblastic leukemia that would be difficult to identify using traditional techniques (15).

Based on this background, we hypothesized that a discovery effort based upon ASE could reveal novel points of fragility in the most resistant tumor cells, the GSCs. GSCs maintained in serum-free conditions maintain both genetic and transcriptional signatures found in the tumors from which they were derived, while removing the contaminating non-transformed cells that complicate genetic discovery. Here, we interrogated ASE in 43 patient-derived GSCs and compare the frequency of ASE to normal tissues to reveal novel dysregulated molecular targets that promote drug resistance and confer therapeutic vulnerabilities.

## RESULTS

### Glioblastoma target gene discovery leveraging recurrent allele-specific expression

To discover genes with ASE, we implemented a statistical model to estimate allele imbalance for every gene (Figure 1A). Our systematic methodology utilizes RNA-seq reads overlapping heterozygous sites and reduces false positives by accounting for technical sources of variation, including sequencing errors, genotyping errors, and overdispersion of RNA-seq read counts. We quantified ASE with RNA allelic imbalance (a_RNA_), which is the difference between the reference allele proportion and the expected value of 0.5. We applied our model to RNA-seq data from 43 patient-derived GSCs representing a diverse population of patients (age, biologic sex, etc.) and tumor features (genetics, transcriptional subgroups, etc.) that we had previously reported (26). We identified 5,808 genes with significant ASE in at least one GSC under a false discovery rate (FDR) of 10% (Supplemental Table S1). ASE was restricted to a single sample for most genes (3,948 out of 5,808). However, 1860 genes showed ASE in 2 or more GSCs and 298 genes showed highly recurrent ASE that was present in 5 or more GSCs (Figure 1B).

**Figure 1:**
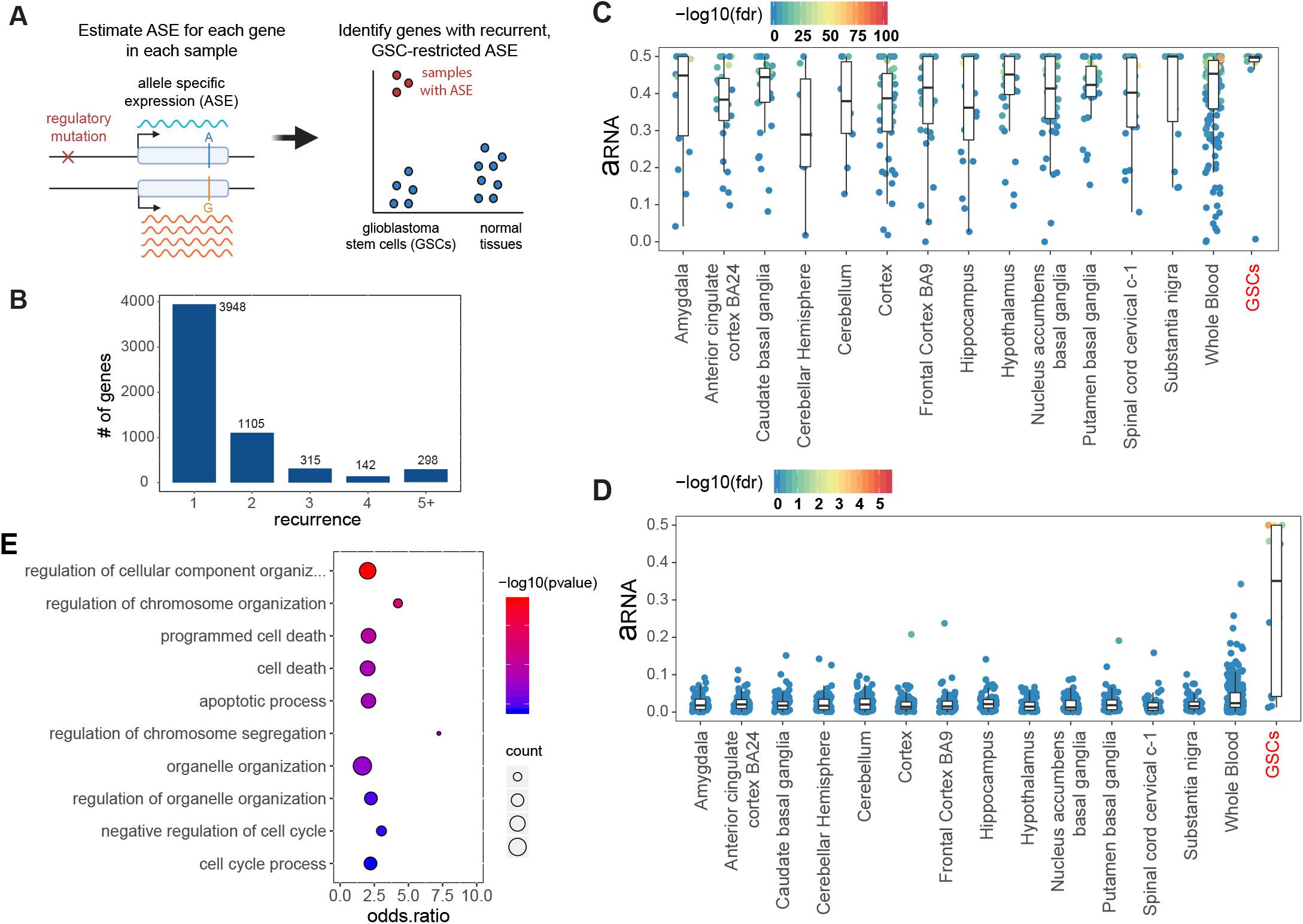
Discovery of genes with recurrent allele-specific expression in glioblastoma stem cells. **A**) Schematic of approach. Allele-specific expression (ASE) is the higher expression of one allele of a gene compared to the other allele and can be used the detect the effects of cis-regulatory mutations. ASE is detectable at genes that contain heterozygous sites. We identify genes that exhibit ASE in glioblastoma stem cells (GSCs) more frequently than in normal tissues. **B**) Recurrence of ASE in GSCs. The histogram indicates the number of GSCs with ASE (FDR corrected p-value ≤ 10%) across 42 patient-derived glioblastoma cell lines, for genes that have ASE in at least one sample. **C**) Estimated ASE (a_RNA_) for GSCs and normal brain and whole-blood samples from GTEx for a known imprinted gene, *H19.* Each point is a sample. Points are colored based on the significance of allele-specific expression (likelihood ratio test). **D**) Estimated ASE for *RHOB*. **E)** Gene ontology analysis of 118 genes that are enriched for ASE in GSCs compared to normal samples.

To discover genes that showed recurrent ASE specific to GSCs relative to normal tissues, we compared the frequency of ASE for each gene in GSCs to both normal whole blood and normal brain samples from the Genotype Tissue Expression (GTEx) project using Fisher’s Exact Test. Whole blood has, by far, the largest number of samples in GTEx so we used it as the initial reference tissue; however, we performed subsequent comparisons with ASE in 13 different brain tissues to account for tissue-specific imprinting, as described below. Under an FDR of 10%, 118 genes displayed significantly enriched ASE in GSCs (Supplemental Table S2). To illustrate the power of this approach, we examined the ASE patterns of the non-coding RNA gene *H19*, which is maternally imprinted (27), and *RHOB* (Ras Homolog Family Member B), which is important for glioblastoma tumorigenesis (28,29). As expected for an imprinted gene, *H19* showed ASE in almost all normal and tumor samples (Figure 1C), however ASE of *RHOB* was restricted to GSCs (Figure 1D).

Recurrent ASE in cancer genomes can be caused by frequent copy number alternations (CNAs) or loss-of-heterozygosity (LOH). In the presence of large CNAs, many genes are expected to exhibit ASE, but these genes would be clustered into the genomic regions that undergo frequent CNAs. However, in our dataset, recurrent ASE genes were distributed throughout the genome, suggesting that their dysregulation was not due to large-scale CNAs, but was instead caused by localized cis-regulatory mutations or epigenetic changes (Supplemental Figure S1A).

To examine the biological function of the 118 genes with recurrent ASE in GSCs, we performed Gene Ontology (GO) enrichment analysis, revealing over-representation of recurrent ASE genes involved in regulation of cell cycle and apoptosis (FDR ≤ 5%). 28 genes with recurrent ASE were associated with the Biological Process GO category “programmed cell death”, whereas only 13 genes were expected by chance (Figure 1E). These genes included the kinase *IP6K2* (Inositol hexakisphosphate kinase 2) (30,31), which exhibited a marked enrichment of ASE in GSCs compared to both whole-blood (FET p-value 0.001; FDR-adjusted p-value 0.06) and brain tissues from GTEx (Supplemental Figure S1B).

### Allele-specific gene expression is associated with H3K27ac marks at distal regulatory elements

Cis-acting pathogenic variants impact gene expression by disrupting regulatory sequences, such as enhancers (18,32). To illuminate the cis-acting mechanisms that underlie ASE in GSCs, we tested whether the expression of the 118 ASE genes was associated with the activity of nearby regulatory elements. We identified putative regulatory sequences within 100-kb of the ASE gene promoters using histone H3 lysine 27 acetyl (H3K27ac) chromatin immunoprecipitation followed by deep sequencing (ChIP-seq) data that we previously generated for the GSCs (26). We divided the genome into 1-kb genomic bins and labeled bins that overlapped H3K27ac peaks in at least one GSC as putative cis-regulatory elements (CREs). To connect CREs with genes, we correlated normalized H3K27ac levels with gene expression, focusing on distal and intronic elements, which exhibit a greater specificity and wider range of activity across cancer samples compared to promoters (33). Of the 118 ASE genes, 56 had a significant Spearman’s rank correlation with the activity of a distal CRE (FDR ≤ 5%). In many cases, a single gene was associated with the activity of multiple CREs, such that 227 CRE bins were associated with the expression of these 56 genes (Supplemental Table S4). Thus, gene expression of many ASE genes was associated with activity levels of distal regulatory elements, as measured by H3K27ac, suggesting that cis-acting mutations within these CREs are a plausible and potentially common mechanism for the dysregulation of these genes in glioblastoma.

One example of an ASE gene that is associated with multiple CREs is *NOTCH1*. ASE of *NOTCH1* was enriched in GSCs compared to both whole-blood (Fisher’s Exact Test p-value 5.09e-5; FDR-adjusted p-value 0.0094) and brain samples and we observed extreme biases in reference and alternate allele proportions at all heterozygous sites (Figures 2A and B). *NOTCH1* gene expression correlated with H3K27ac levels of 29 nearby CREs (Figure 2C; Supplemental Table S4). To confirm that these CREs were more strongly associated with *NOTCH1* expression than expected by chance, we created an empirical null distribution for each CRE by correlating H3K27ac levels with the expression of 1000 randomly selected genes. Using these null distributions, 22 out 29 CREs remained significantly correlated with *NOTCH1* gene expression (empirical p-value ≤ 5%) (Supplemental Table S5). These regions may contain cis-regulatory non-coding mutations that lead to *NOTCH1* gene dysregulation and may be excellent targets for future regulatory screens dissecting the regulation of *NOTCH1* expression in GSCs.

**Figure 2:**
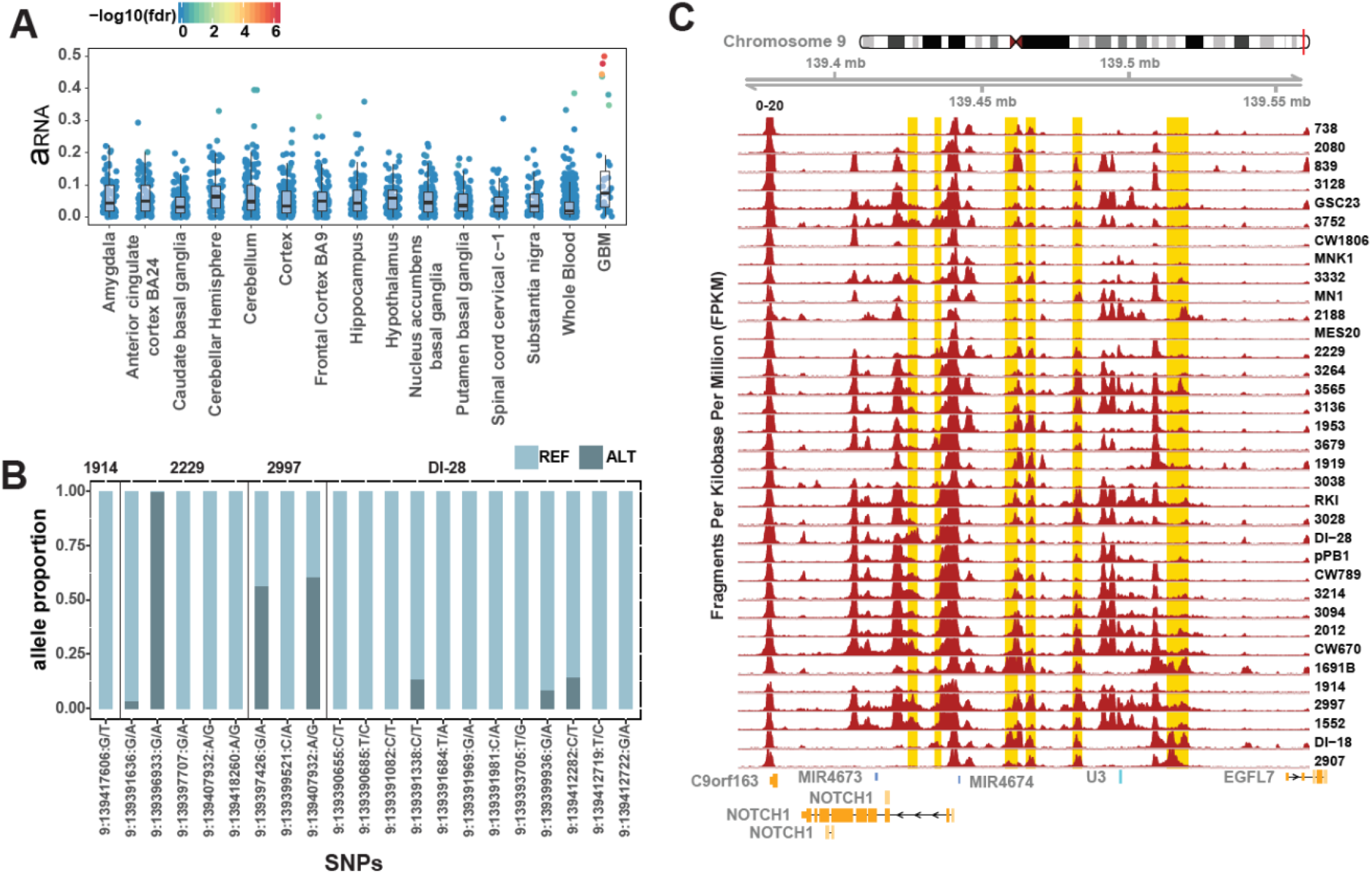
*NOTCH1* exhibits recurrent allele-specific expression and association with multiple cis-regulatory elements. **A)** Allele-specific expression (ASE) estimates (a_RNA_) of *NOTCH1* in glioblastoma stem cells and normal brain and whole-blood samples from GTEx. **B)** The proportion of *NOTCH1* RNA-seq reads from the reference and alternate alleles at heterozygous sites in 4 GSCs with significant *NOTCH1* ASE. **C)** The H3K27ac profile around *NOTCH1*. Samples are arranged in increasing order of gene expression with the highest expression samples at the bottom. Putative cis-regulatory elements that are correlated with *NOTCH1* gene expression (empirical p-value ≤ 0.05) are highlighted in gold.

### Promoter methylation is associated with the expression of ASE genes

Aberrant DNA methylation of gene promoters is associated widespread gene expression changes and chemotherapy resistance in brain tumors. In gliomas, *MGMT* CpG-rich promoter methylation is associated with decreased expression and improved response to temozolomide (TMZ) treatment (34). To determine whether genes with recurrent ASE were associated with aberrant DNA methylation, we analyzed the CpG methylome of the GSCs (26). We estimated the promoter methylation of the 118 ASE genes (β_promoter_) and computed Spearman’s rank correlations with normalized gene expression. We identified 30 genes that displayed correlation between promoter methylation and gene expression at FDR ≤ 10% (Supplemental Table S6), of which 16 were correlated with the H3K27ac levels of nearby CREs. Thus, 70 of the 118 ASE genes were associated with promoter methylation, CRE activity or both, suggesting possible mechanisms for their dysregulation.

### *SLFN11* promoter methylation associates with its gene expression

One of the 30 genes with correlated gene expression and promoter methylation was *SLFN11* (rho=-0.68, FDR corrected p-value = 0.0001) (Figures 3A-C, Supplemental Table S5). *SLFN11* is notable because it inhibits DNA replication and promotes cell death in response to DNA damage (35,36). Loss of *SLFN11* causes resistance to poly ADP ribose polymerase (PARP) inhibitors in small cell lung cancer, suggesting that it may be an important marker of chemotherapy resistance (37). ASE of *SLFN11* was highly enriched in GSCs with 4 out of 10 testable GSCs exhibiting ASE, compared to 0 out of 159 testable normal whole blood tissues from GTEx (FET p-value 6.4e-6; FDR-adjusted p-value 2.2e-3) (Supplemental Table S2). Although this gene showed enrichment of ASE in GSCs compared to whole blood samples, we detected ASE in small number of normal brain tissue samples (4 out of 224) (Figure 3C). This suggests that rare germline variants or somatic events, such as mutations or DNA methylation, may affect *SLFN11* expression in some phenotypically normal individuals.

**Figure 3:**
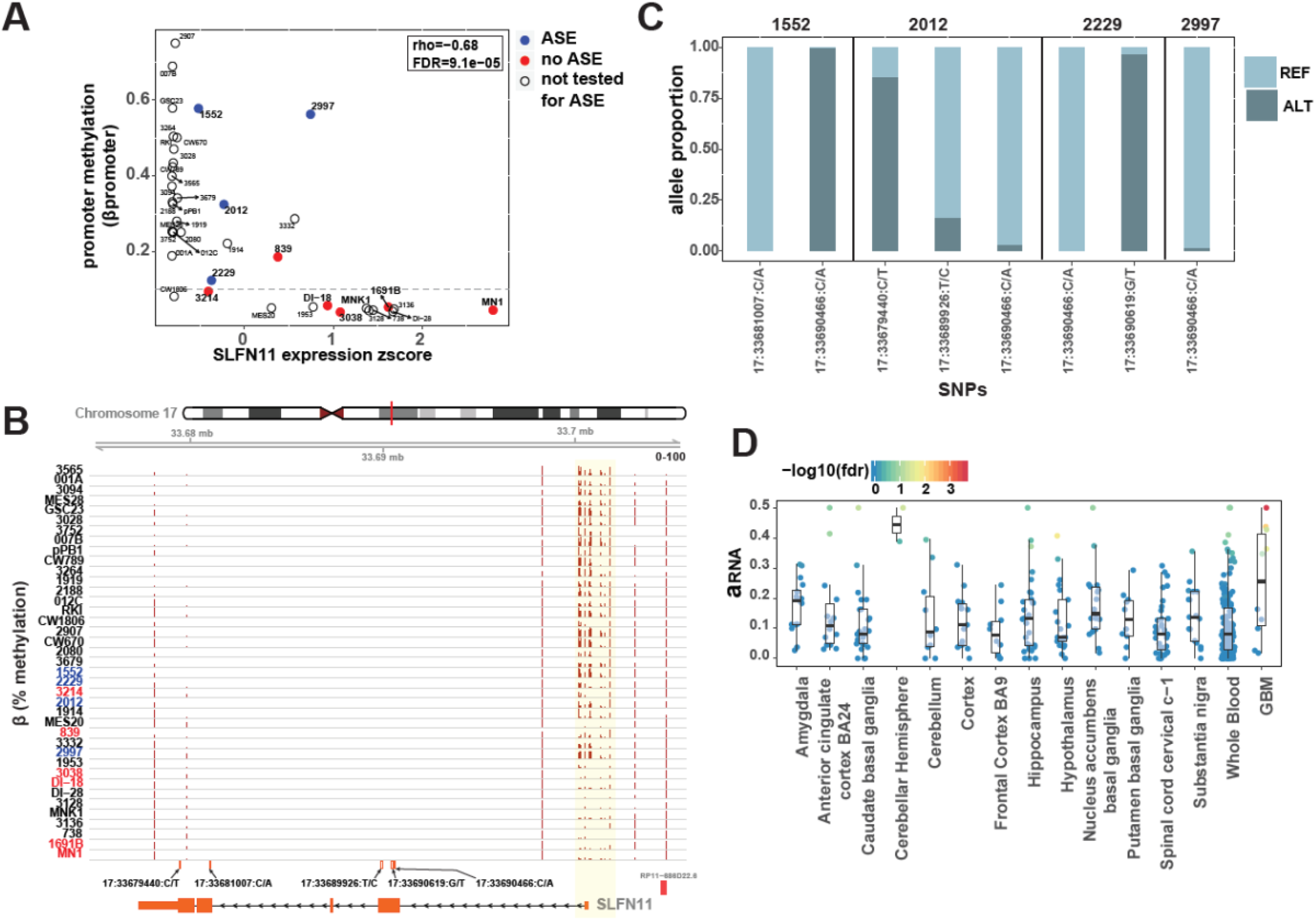
*SLFN11* gene expression and allele-specific expression are correlated with promoter methylation. **A)** Scatterplot showing correlation between gene expression and mean promoter methylation (β_promoter_) in glioblastoma stem cells (GSCs). Samples where *SLFN11* allele-specific expression (ASE) could be estimated are colored blue if they have significant ASE and red if they do not have significant ASE. **B)** CpG methylation around promoter regions of *SLFN11*. Samples are arranged in increasing order of gene expression. **C)** Reference and alternate allele proportions of RNA-seq reads at heterozygous sites for the 4 GSCs with significant *SLFN11* ASE. **D)** Estimated ASE for *SLFN11* in GSCs compared to normal brain and whole-blood samples from GTEx.

Based on the promoter methylation and gene expression of *SLFN11*, GSCs can be divided in 3 distinct classes: 1) GSCs with high methylation and low expression; 2) GSCs with hemi-methylation and intermediate expression; and 3) GSCs with low methylation and high expression (Figure 3A). Four GSC samples with ASE of *SLFN11* had detectable ASE within the class of hemi-methylated samples, in which one allele was silenced and the other was expressed. However, samples like GSCs 2907 and 007B had high promoter methylation levels (>60%) and very low expression of *SLFN11* (Figures 3A and B). Under these circumstances, genes would not be found by ASE because the expression of both alleles is reduced. These results demonstrate that low expression of *SLFN11* in GSCs is associated with increased promoter methylation and the samples with detectable ASE are consistent with hemi-methylation.

### *SLFN11* augments chemotherapy resistance in GSCs

*SLFN11* regulates cellular responses to DNA damaging agents (35,36). Therefore, we hypothesized that *SLFN11* expression in GSCs would be associated with chemotherapeutic resistance to an alkylating agent, TMZ, and a PARP inhibitor, olaparib. To test this hypothesis, we utilized four patient-derived GSCs: two with high *SLFN11* expression and no evidence for ASE (839 and MNK1) and two with low *SLFN11* expression and strong ASE (2012 and 1552). We confirmed both *SLFN11* mRNA and SLFN11 protein abundance by RT-PCR (Figure 4A) and immunoblot (Figure 4B). We treated cells with drug concentrations ranging from 0 to 1000 μM for TMZ and 0 to 50 μM for olaparib, then measured drug effects on cell survival to generate a concentration-response matrix where an effect of 100% corresponds to complete killing of all cells and 0% corresponds to no difference in cell survival (Figures 4C and D; Supplemental Figure S1). We estimated synergy between the two drugs using SynergyFinder 2.0 (38). Two GSCs with low expression and ASE of *SLFN11* had reduced responses to drug treatment compared the GSCs with high expression of *SLFN11*. The maximum response for both ASE GSCs ranged from 40-50%, whereas the GSCs with high expression of *SLFN11* had responses ranging from ~70-80% (Figure 4D). The two ASE GSCs also had lower drug synergy scores for the combination of the drugs (Figure 4E).

**Figure 4:**
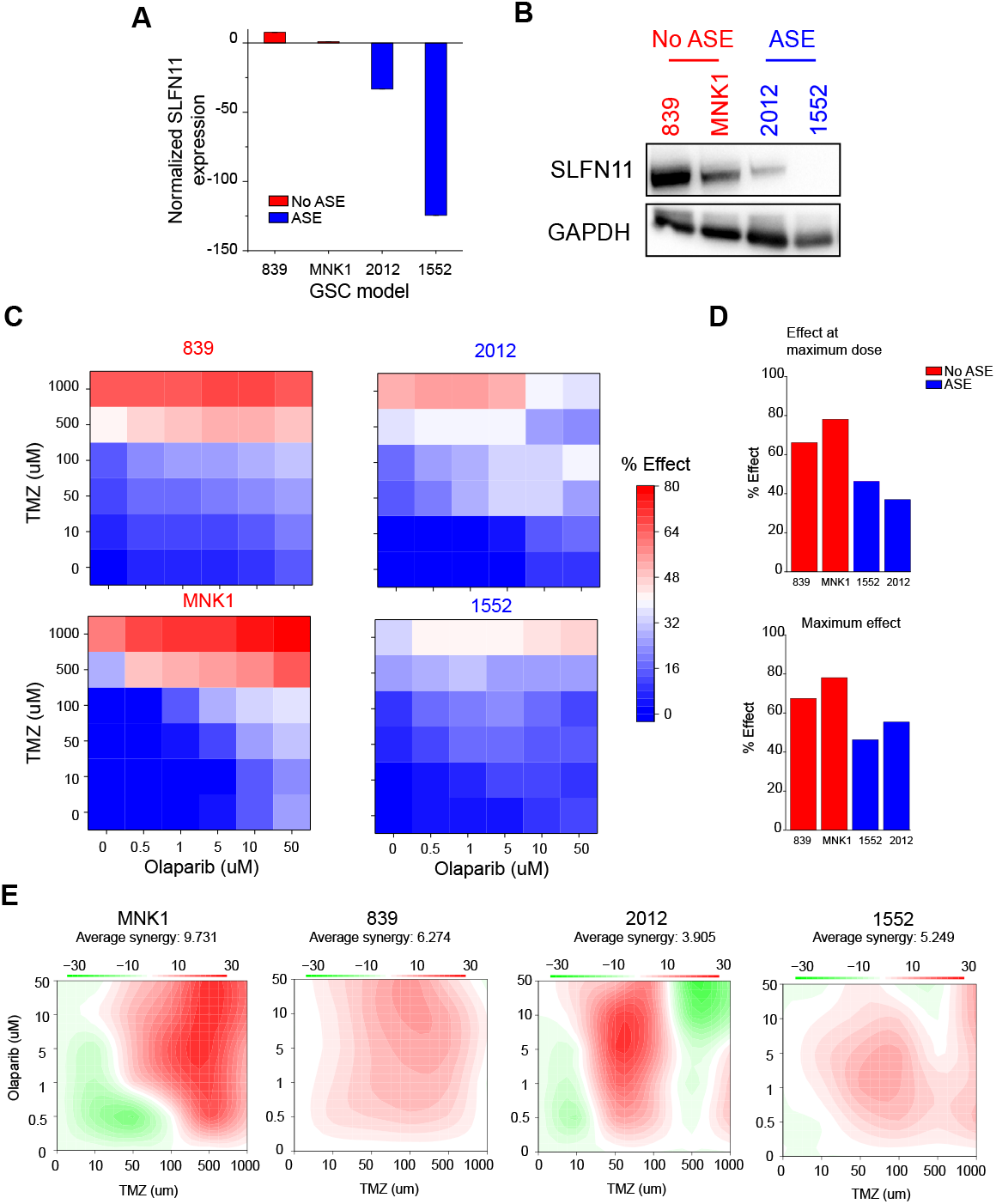
*SLFN11* expression is associated with chemotherapeutic resistance in GSCs. **A)** qPCR of *SLFN11* expression in GSCs without ASE of *SLFN11* GSCs 839 and MNK1 (red) compared to GSCs with ASE of *SLFN11* 2012 and 1552 (blue). Gene expression is plotted as 2^−ΔΔCt^, with *SLFN11* expression for each sample normalized to *GAPDH* expression and then compared to *MNK1*. Values <1 were transformed by taking the negative inverse. Data are represented as mean +/− standard deviation. **B)** Western blot of the same non-ASE (red) and ASE (blue) GSCs for SLFN11 and GAPDH expression. **C)** Cell viability of each GSC (left: non-ASE; right: ASE) following treatment with temozolomide (TMZ) and olaparib. Cell viability relative to DMSO control is annotated on a blue-white-red scale with blue indicating high viability, or minimal drug effect, and red indicating low viability, or strong drug effect. Doses are scaled categorically on the x-axis (olaparib) and y axis (TMZ). **D)** Top: effect of the maximum combinatorial drug dose on cell viability for non-ASE (red) and ASE (blue) GSCs. Bottom: maximal effect achieved at any dose for each model. Percent effect on reduction of cell viability is measured on the Y-axis. **E)** Synergy of TMZ and olaparib for each model, with red indicating high synergy and green indicating antagonism.

To directly test whether *SLFN11* expression affects chemotherapeutic drug sensitivity in GSCs, we also performed knockdown (KD) and overexpression (OE) experiments. Specifically, we performed KD of *SLFN11* in MNK1 GSCs, which have high baseline expression of *SLFN11* (Figure 5A), and *SLFN11* OE in 2012 GSCs, which have low baseline expression of *SLFN11* (Figure 5B). After confirming the expression of *SLFN11* in MNK1 and 2012 GSCs, we measured cell survival in response to treatment with TMZ and olaparib. *SLFN11* KD in MNK1 cells reduced cell death in response to drug treatment, and conversely *SLFN11* OE in 2012 cells increased cell death in response to both drugs (Figures 5C and D). Synergistic drug response to TMZ and olaparib treatment decreased upon *SLFN11* KD and increased with OE (Figure 5E). Thus, *SLFN11* expression is causally linked to sensitivity to the chemotherapeutic drugs TMZ and olaparib.

**Figure 5:**
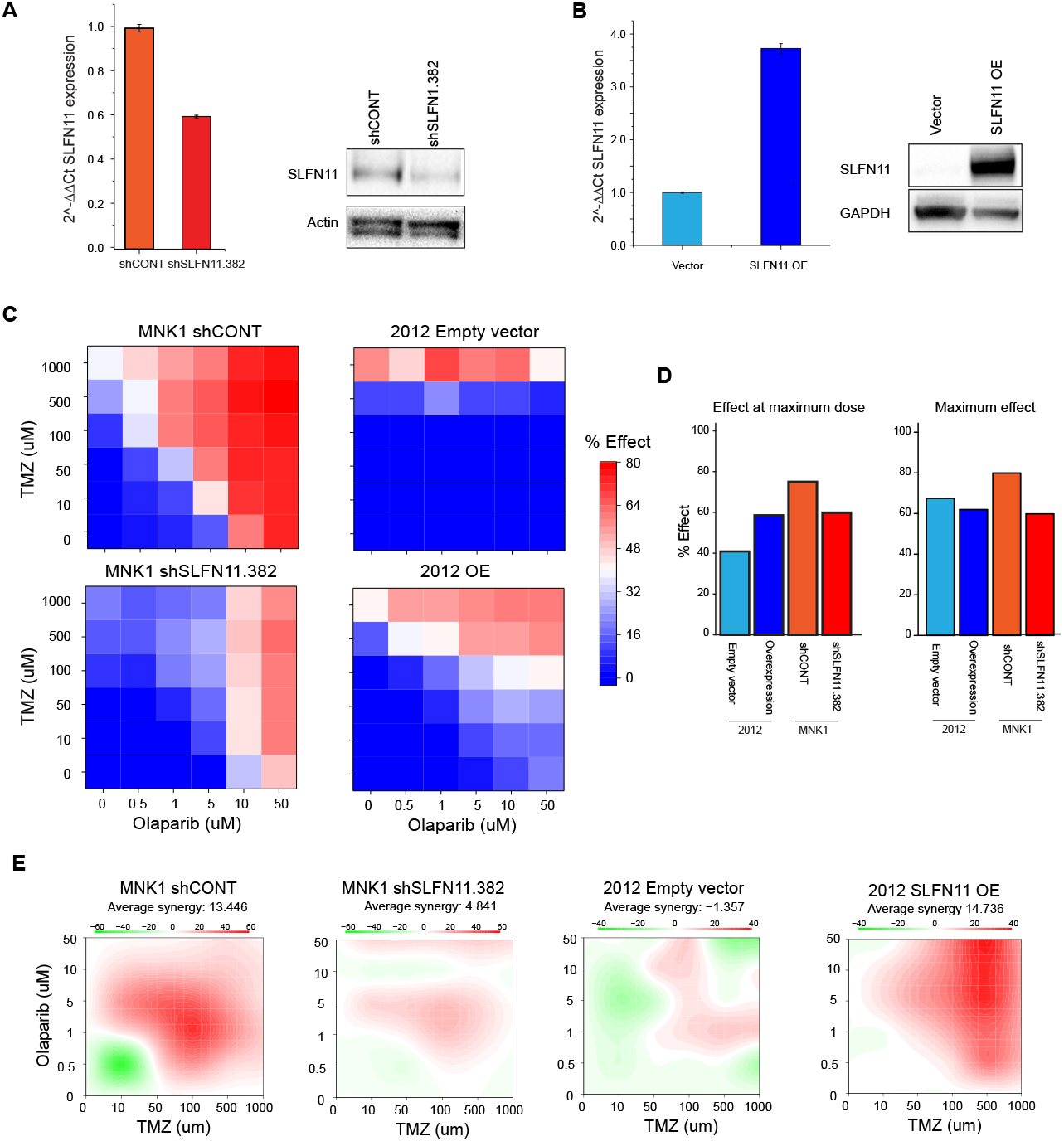
Knockdown and overexpression of *SLFN11* modulates chemotherapeutic resistance in GSCs. **A)** Left: qPCR of SLFN11 expression in MNK1 GSCs transduced with a non-targeting control shRNA (shCONT) or a shRNA targeting *SLFN11* (shSLFN11.382). Gene expression is plotted as 2^−ΔΔCt^ normalized to Actin expression. Right: Western blot of the same samples for SLFN11 and Actin expression. An un-spliced version of the Western blot is provided in Supplemental Figure S3C. **B)** Left: qPCR of *SLFN11* expression in GSC 2012 transduced with an empty vector or a *SLFN11* overexpression vector (OE). Gene expression is plotted as 2^−ΔΔCt^ normalized to *GAPDH* expression. Right: Western blot of the same samples for *SLFN11* and *GAPDH* expression. **C)** Cell viability of MNK1 GSCs transduced with shCONT (top left) or shSLFN11.382 (bottom left) and of 2012 GSCs transduced with an empty vector (top right) or OE vector (bottom right) following treatment with temozolomide (TMZ) and olaparib. Cell viability relative to DMSO control is annotated on a blue-white-red scale with blue indicating high viability, or minimal drug effect, and red indicating low viability, or strong drug effect. Doses are scaled categorically on the x-axis (olaparib) and y axis (TMZ). **D)** Left: effect of the maximum combinatorial drug dose on cell viability for each comparison. Right: maximal effect achieved at any dose for each model. Percent effect on reduction of cell viability is measured on the Y-axis. **E)** Synergy of TMZ and olaparib for each model, with red indicating high synergy and green indicating antagonism. Error bars in panels A and B are mean +/− standard deviation.

### GSCs with low expression of *SLFN11* are sensitive to Zika virus

*SLFN11* may be involved in cellular response to viral infection. *SLFN11* is upregulated following virus-induced type I interferon response and restricts flavivirus replication in human tumor cell lines (39-41). Oncolytic viruses that infect the central nervous system can be leveraged to treat brain tumors (42,43). We recently demonstrated that Zika virus, which is a member of the flavivirus genus of RNA viruses, preferentially infects and kills GSCs compared to differentiated tumor cells and normal neuronal cells (44). Based on this background, we hypothesized that tumor cells with promoter methylation of *SLFN11* would be more susceptible to oncolytic destruction by Zika because they are unable to increase *SLFN11* expression in response to interferon stimulation (44). To test this hypothesis, we examined the association between *SLFN11* gene expression and type I interferon response by analyzing gene expression data from 669 glioblastoma tumors obtained from The Cancer Genome Atlas (TCGA) (45). To quantify interferon response in each sample, we computed a single-sample gene set enrichment analysis (ssGSEA) score using the interferon alpha response hallmark gene set (46-48). Glioblastoma had higher gene expression of *SLFN11* compared to grade II and III gliomas (Figure 6A), and *SLFN11* gene expression correlated with the interferon alpha (IFNα) ssGSEA score (R=0.4, p-value <2.2e-16), consistent with induction of *SLFN11* expression in response to interferon signaling (Figure 6B). Glioblastomas with higher expression of *SLFN11* had gene set enrichment for activation of immune response, complement activation, defense response to virus, and response to interferon alpha compared to tumors with lower *SLFN11* expression (Figure 6C). Enrichment of these immune pathways likely reflects the constitutive activation of autocrine interferon signaling, which might facilitate immune escape for tumors (49).

**Figure 6:**
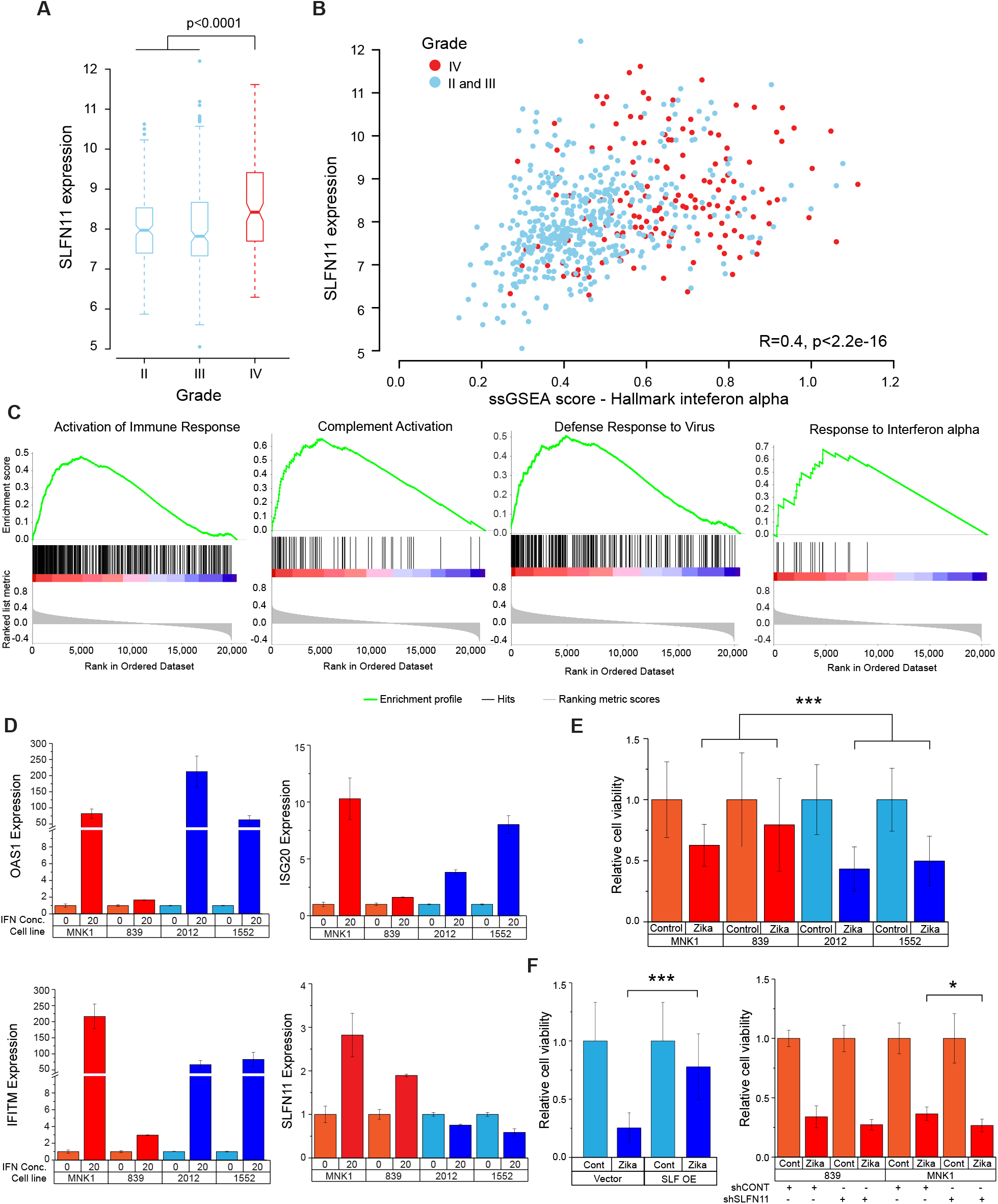
*SLFN11* expression affects sensitivity of GSCs to Zika virus. **A)** Box plot of *SLFN11* expression in The Cancer Genome Atlas database comparing grade II, III and IV gliomas. Boxes are notched at the median and extend from the first to third quartile, with whiskers extending from 5%-95%. **B)** Correlation of *SLFN11* expression with ssGSEA score for the Hallmark IFNα geneset by tumor. Grade II and III tumors are in blue and glioblastoma tumors are in red. **C)** GSEA plot of *SLFN11* expression correlation with selected immune signatures from GO biological process or Reactome datasets. **D)** qPCR expression of IFNα-induced genes *OAS1*, *IFITM*, *ISG20* and *SLFN11* following 8 hours of treatment with 20 ng/mL of IFNa. Gene expression is plotted as 2^−ΔΔCt^ normalized to *GAPDH* expression. **E)** Cell viability following incubation with Zika virus vs empty control virus. Non-ASE samples are in blue, ASE samples are in red. Data were compared using two-way ANOVA for cell line and ASE status. **F)** Cell viability following incubation with Zika virus vs control in 2012 GSCs transduced with a SLFN11 overexpression (OE) vector (left) or empty vector (right). Student’s t test was used to test for differences in expression **G)** Cell viability following incubation with Zika virus (dark blue) vs control in *SLFN11* KD vs control 839 (left) or MNK1 (right) GSCs. Mean expression was compared using Student’s t test. Data are represented as mean +/− standard deviation. * <0.05; ** <0.01, *** <0.0001.

To more directly test whether interferon signaling induces *SLFN11* gene expression, we treated GSCs with IFNα and measured the expression of *SLFN11* and type 1 immune response genes, *OAS1*, *ISG20*, and *IFITM* 8 hours after treatment. All four GSCs increased expression of *OAS1*, *ISG20*, and *IFITM* following IFNα treatment, although the level of induction in the GSC 839 was modest (2-3-fold) compared to the other GSCs (Figure 6B). IFNα treatment also increased *SLFN11* gene expression in MNK1 and 839 GSCs, which have high baseline expression of *SLFN11* and no ASE. However, IFNα treatment did not increase expression of *SLFN11* in 2012 and 1552 GSCs, both of which showed ASE of *SLFN11* and high promoter methylation (>25% β_promoter_). These results suggest that DNA methylation of the *SLFN11* promoter blocks upregulation by interferon signaling (Figure 6D).

To determine if GSCs with promoter methylation and low expression of *SLFN11* were more susceptible to killing by Zika, we treated the same four GSCs with Zika or saline control (Figure 6E). GSCs with promoter methylation of *SLFN11* (2012 and 1552) displayed decreased viability following Zika infection compared to GSCs without promoter methylation (839 and MNK1). To directly test whether *SLFN11* expression affects susceptibility to Zika, we performed *SLFN11* OE in 2012 GSCs (which have low baseline expression of *SLFN11*) and *SLFN11* KD in the 839 and MNK1 GSCs (which have high baseline expression of *SLFN11*). *SLFN11* OE decreased susceptibility to Zika (Figure 6F), while KD increased susceptibility to Zika (Figure 6G). These results demonstrate that *SLFN11* expression is critical for immune response and cell viability following Zika infection.

## DISCUSSION

ASE analysis of tumor genomes is a new approach that can discover biomarkers and therapeutic targets by illuminating genes that are recurrently dysregulated, even when the identity of the mutational events that drive this dysregulation is unknown (50,51). In this study, we utilized ASE to identify recurrently dysregulated genes in GSCs derived from patient surgical specimens. Using this approach, we discovered 118 candidate disease genes that were recurrently dysregulated in GSCs, but not in normal tissues. Our ASE analysis revealed genes with an established role in tumor biology. For example, *IP6K2*, a pro-apoptotic protein kinase showed ASE almost exclusively in GSCs. *IP6K2* selectively binds to HSP90, which decreases its catalytic activity and inhibits apoptosis (31). Disruption of this interaction by cisplatin and novobiocin, chemotherapeutic compounds that bind to the C-terminus of HSP90, restores its catalytic function and promote apoptosis (52). Furthermore, knockdown of *IP6K2* in colorectal cancer cells has been demonstrated to selectively impair p53-mediated apoptosis, instead favoring cell-cycle arrest (53). These observations from previous studies suggest that *IP6K2* may be an important tumor suppressor in glioblastoma.

*NOTCH1* also exhibited ASE that was specific to GSCs. *NOTCH1* regulates neural stem cell fate during neurogenesis and high expression of *NOTCH1* has been reported in many high-grade gliomas (54-57). Notch1 signaling promotes invasion, self-renewal, and growth of GSCs (58-60); *NOTCH1*-KD suppresses cell proliferation and induces apoptosis (61). Furthermore, inhibition of the Notch1 signaling pathway sensitized tumor cells to apoptosis induced by ionizing radiation, the death ligand TRAIL (tumor necrosis factor-related apoptosis-inducing ligand), or the Bcl-2/Bcl-XL inhibitor ABT-737 (62). These studies suggest that *NOTCH1* may help maintain the stem-cell like behavior of GSCs and promote tumor progression. The multiple CREs correlated with *NOTCH1* expression are potentially excellent targets for subsequent studies and cis regulatory screens.

Recurrent ASE of *SLFN11* is an important finding because this gene has recently emerged as a biomarker of drug sensitivity in cancer (63). We demonstrate that in GSCs, *SLFN11* gene expression is associated with DNA methylation of its promoter and its expression is required for the anti-tumor activities of the DNA alkylating agent temozolomide (TMZ) and the replication-inhibitor olaparib. The current standard-of-care for glioblastoma patients is maximum safe surgical resection followed by concurrent TMZ and radiation therapy (64). Similar to *MGMT* promoter DNA methylation, *SLFN11* promoter methylation may be a biomarker that predicts the efficacy of DNA damaging agents, such as TMZ and olaparib. In addition, promoter methylation and reduced expression of *SLFN11* may reflect evolution of resistance to TMZ within tumor cells. In GSCs, *SLFN11* gene expression was regulated by type 1 interferons and frequently upregulated in high-grade tumors, alongside constitutive activation of autocrine interferon signaling that facilitates immune evasion of GBM cancer cells (49). However, in the presence of promoter CpG methylation, *SLFN11* is unresponsive to IFNα cytokine treatment, rendering GSCs vulnerable to killing by oncolytic viruses, such as Zika (44). Thus, tumors refractory to DNA damaging agents may be more amenable to treatment with genetically modified viruses. As GSCs with low *SLFN11* expression are susceptible to Zika, but resistant to chemotherapy and vice-versa, the combination of oncolytic viruses and chemotherapy may be a powerful treatment approach (Figure 7).

**Figure 7:**
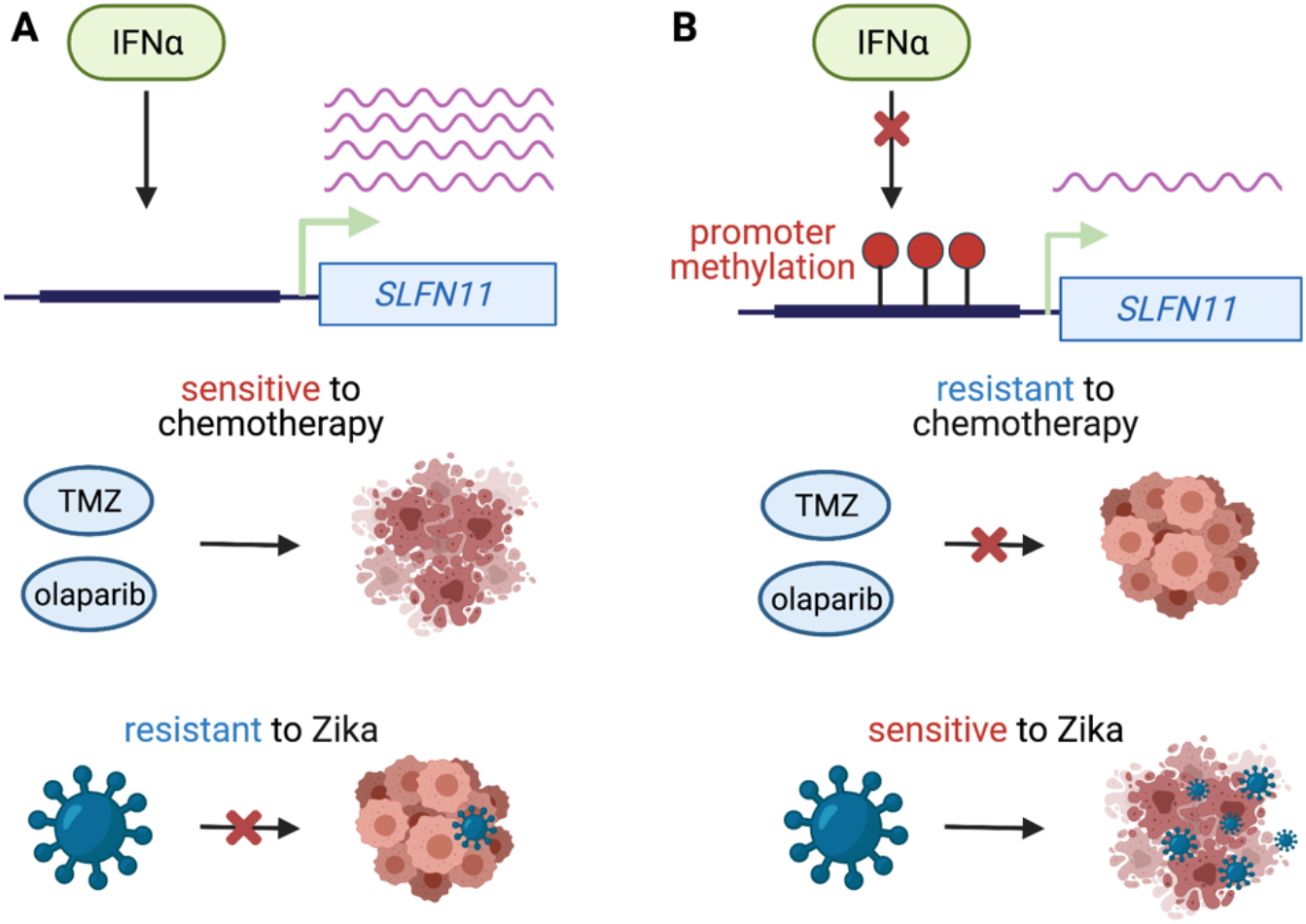
Model of *SLFN11* expression and therapeutic response. **A)** Interferon alpha increases *SLFN11* expression when its promoter is unmethylated. Glioma stem cells (GSCs) with high *SLFN11* expression are sensitive to temozolomide (TMZ) and olaparib because they undergo apoptosis in response to DNA damage. However, GSCs with high *SLFN11* expression are resistant to Zika virus because *SLFN11* restricts flavivirus replication. **B)** Interferon alpha fails to induce *SLFN11* expression when its promoter is methylated. Glioma stem cells with low expression of *SLFN11* are resistant to chemotherapy, but sensitive to Zika virus.

## METHODS

### Derivation and maintenance of GSCs

Glioblastoma samples were obtained from surgical resection from patients at Duke University or Case Western Reserve University with informed consent in accordance with the Cleveland Clinic Institutional Review Board-approved protocol 090401. Prior to use, all samples were reviewed and verified by a neuropathologist. All patient studies were conducted in accordance with the Declaration of Helsinki. GSC23 was acquired via a material transfer agreement from The University of Texas MD Anderson Cancer Center (Houston, TX). GSCs were cultured in Neurobasal media (Invitrogen) supplemented with B27 without vitamin A (Invitrogen), EGF, and bFGF (20 ng/mL each; R&D Systems), sodium pyruvate, and Glutamax. Short tandem repeat analyses were performed to authenticate the identity of each tumor model used in this article on at least yearly basis. Cells were stored at −160°C when not being cultured. To minimize cell culture–based artifacts, patient-derived xenografts were produced and propagated as a renewable source of tumor cells for study.

### Variant calling

Exome-seq reads from GSCs were aligned to the GRCh37 (hg19) assembly of the reference genome using BWA-MEM with default parameters (65). Mapped reads were filtered for mapping quality score ≥ 30 and duplicate reads were removed using samtools (1.9) (66). Genotypes were generated for each individual using GATK’s *HaplotypeCaller* and jointly processed using the *GenotypeGVCFs* function in GATK (4.1.1). Following genotyping, single nucleotide variants (SNVs) were extracted and filtered using Variant Quality Score Recalibration (VQSR) in GATK (4.1.1).

### RNA-seq alignment and processing

RNA-seq reads were aligned end-to-end to the GRCh37 (hg19) assembly of the reference genome using STAR (2.5.3a) (67). Mapped reads were filtered using mapping quality score ≥ 20 and duplicate reads were removed using samtools (1.9) (66). Read counts for GENCODE (v28) genes were computed using *FeatureCounts* (1.6.3) (68) and Fragments Per Kilobase Per Million (FPKM) values were estimated using DESeq2 (1.14.1) (69). For downstream analysis, the FPKM values were quantile normalized, and then converted to z-scores by mean-centering and standardizing across samples.

### Estimating allele-specific expression

RNA-seq read alignments were corrected for mapping bias and allele-specific read counts at all heterozygous positions were collected using WASP (70). The filtered read counts obtained from WASP were used for modelling allele-specific expression (ASE) for each gene.

Misclassification of heterozygous sites can occur due to incorrect genotypes. To control for genotyping errors, we calculate the error rate (*ϵ*_*S*_) directly from genotype quality scores (GQ) from GATK:

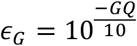

Incorrect allele-specific read counts can be caused by sequencing errors in reads. To control for sequencing errors, we approximate the sequencing error rate, *ϵ*_*S*_, using the count of “other” reads which do not match reference or alternate allele, *X_0_*. To account for the fact that only 2/3 of sequencing errors will be observed in the “other” reads (the other 1/3 will match the alternate or reference allele), we scale the error rate estimate, by 3/2:

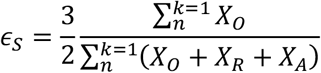

We assume a heterozygous site is equally likely to be misclassified as homozygous reference or alternate. Conditional upon a genotyping error having occurred, we define the likelihood at site *i* as:

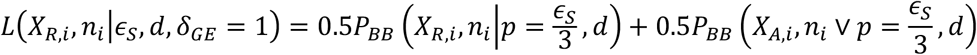

and conditional on no genotyping error the likelihood is:

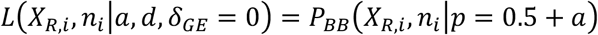

where *P*_*BB*_ is the Beta Binomial probability distribution function, *n*_*i*_ is the total count of reads matching the reference or alternative sequence (*n*_*i*_ = *X*_*R*,*i*_ + *X*_*A*,*i*_) at heterozygous site *i*, *a* is the allelic imbalance parameter defined over the range [-0.5,0.5], *d* is the dispersion parameter, and *δ*_*GE*_ is an indicator variable that is 1 when a genotyping error has occurred and 0 otherwise. We estimate *d* by maximum likelihood over all heterozygous sites overlapping exons (setting *a* to 0). Finally, a single gene might contain multiple heterozygous sites which need to be combined to estimate the allele-imbalance for a gene. We define the likelihood of the read counts for the first heterozygous site within a gene as:

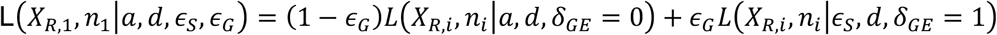

For subsequent heterozygous sites in the same gene, we do not know the phase of the alleles with respect to the first heterozygous site. We assume that the reference and alternative alleles are equally likely to be on the same haplotype as the reference allele at the first site. The combined likelihood of all sites within a gene is then defined as:

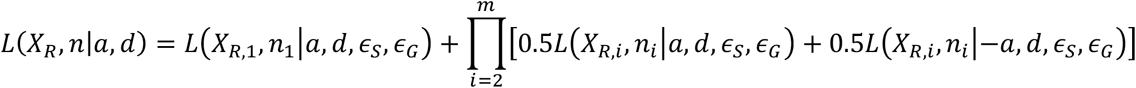

We estimate *a* for each gene by maximum likelihood under the alternative model of allelic imbalance. Then we use a likelihood ratio test to compare the alternative model to the null model of no allelic imbalance (i.e. with *a* fixed to *a* = 0). We correct the p-values from the likelihood ratio test for multiple testing using the Benjamini-Hochberg method. To make it clear when we are referring to allelic imbalance in RNA instead of DNA, we subsequently refer to *a* for RNA-seq reads as *a*_*RNA*_.

### Enrichment compared to a normal whole-blood and brain tissues from GTEx

To generate a reference profile of ASE in normal samples, we obtained RNA-seq data for 369 whole blood and 216 brain samples distributed across 13 brain regions from the GTEx consortium [19] and analyzed this data using our ASE method. To discover genes which were enriched for ASE in GSCs, we compared the frequency of samples with significant ASE for each gene between GSCs and whole blood using a Fisher Exact Test (FET) and adjusted the resulting p-values using the Benjamini-Hochberg procedure. We only analyzed genes which were testable for ASE (i.e., had ≥ 1 heterozygous site with ≥10 reads) in both GSCs and whole blood tissues. For further analysis of ASE frequency for individual genes we also compared estimated allelic imbalance from our model (a_RNA_) between whole blood, 13 brain tissues, and GSCs. Manhattan plots for enriched genes were generated using ggbio (1.30). Gene ontology enrichment analysis for genes showing recurrent ASE in GSCs was carried out using the topGO (2.34)(71).

### Association between DNA methylation and gene expression

We downloaded pre-computed genome-wide methylation data for 43 GSCs from Gene Expression Omnibus (GSE119774) (26). This methylation data was generated using the Illumina Infinium Epic Methylation Array. In this assay, DNA methylation levels at CpG sites are represented by β which is the ratio of the methylated (C) to unmethylated (T) signal. We annotated the CpG probe positions based on GENCODE (v28) genes and computed the mean β values for promoter regions (i.e. 1-kb upstream to 500-bp downstream of annotated transcription start sites) (β_promoter_). To discover ASE genes which may be dysregulated by aberrant DNA methylation, we computed Spearman’s rank correlation between β_promoter_ and normalized gene expression. We corrected correlation p-values for multiple testing using the Benjamini-Hochberg procedure. For this analysis we only considered genes with ≥ 3 CpG probes mapping to their promoter regions.

### Analysis of H3K27ac chromatin immunoprecipitation and sequencing data

We downloaded H3K27ac chromatin immunoprecipitation and sequencing (ChIP-seq) data for 35 GSCs from Gene Expression Omnibus (GSE119755) (26). ChIP-seq reads were aligned to the GRCh37 (hg19) assembly of the reference genome using BWA-MEM with default parameters (65). The mapped reads were filtered using mapping quality score of ≥20 and duplicate reads were removed using samtools (1.9) (66). H3K27ac peaks were called using MACS2 in paired-end mode with custom parameters (--nomodel --extsize 200 --qvalue 0.05) (72). To generate a unified set of test regions, we divided the genome in 1-kb non-overlapping genomic bins and kept the bins which overlapped ChIP-seq peaks in at least 1 GSC. We recounted the reads mapping to these genomic bins using exomeCopy (1.28.0) and calculated fragments per kilobase per million (FPKM) using DESeq2 (1.22.2) (69). FPKM measurements were further quantile normalized and mean-centered for downstream analysis.

The 1-kb genomic bins generated from H3K27ac ChIP-seq peaks were annotated using the ChIPSeeker (1.18.0) package (73). To discover cis-regulatory elements (CREs), we selected all distal intergenic and intronic genomic bins located within 100-kb of promoters (i.e. 1-kb upstream and 500-bp downstream of transcription start sites) of ASE genes. We performed a Spearman’s correlation analysis between normalized coverage for bins and normalized expression for genes to identify CREs. We corrected the p-values for multiple testing using the Benjamini-Hochberg procedure.

### Quantitative RT-PCR

Trizol reagent (Sigma-Aldrich) was used to isolate total cellular RNA from cell pellets and qScript cDNA Synthesis Kit (Quanta BioSciences) was used for reverse transcription. Quantitative real-time PCR was performed using SYBR-Green PCR Master Mix (Thermo Fisher Scientific) on an Applied Biosystems 7900HT cycler.

### Western blotting

Cells were collected and lysed in RIPA buffer (50 mM Tris-HCl, pH 7.5; 150 mM NaCl; 0.5% NP-40; 50 mM NaF with protease inhibitors) and incubated on ice for 30 minutes. Lysates were centrifuged at 4°C for 15 minutes at 14,000 rpm, supernatant was collected, and protein concentration was confirmed using a Pierce BCA protein assay kit (Thermo Scientific, cat #23225). Equal amounts of protein samples were mixed with SDS Laemmli loading buffer, boiled for 10 minutes, electrophoresed using NuPAGE Bis-Tris gels and transferred onto PVDF membranes. Membranes were blocked for 1 hour with TBS-T plus 5% non-fat dry milk, then incubated in primary antibodies overnight at 4°C. Blots were washed 3 times for 5 minutes each with TBS-T and then incubated in TBS-T plus 5% milk for 1 hour with appropriate secondary antibodies. Blots were imaged using BioRad Image Lab software and processed using Adobe Illustrator to create the figures. The following antibodies were used for Western blot: SLFN11 (Santa Cruz Biotechnology, cat #SC-374339) and HRP-conjugated GAPDH (Proteintech, cat #HRP-60004).

### Lentiviral transduction

Lentiviral constructs expressing shRNAs directed against *SLFN11* (Sigma TRCN, TRCN0000152057) or a non-targeting control shRNA (TRCN0000231489) with no targets in the human genome were obtained from Sigma-Aldrich. The *SLFN11* expression vector was obtained from VectorBuilder along with an empty vector control with the same lentiviral backbone. 293T cells were used to generate lentiviral particles by co-transfection of packaging vectors psPAX2 and pMD2.G using a standard PEI transfection method in DMEM media plus 1% penicillin/streptinomycin. GSCs were transduced with the lentiviral constructs, and selection was started 48 hours later using 1 μg/mL of puromcyin for 72 hours, at which times cells were assayed for *SLFN11* expression.

### In vitro treatment and cell viability

For *in vitro* cell viability assays, 2000 cells/well for individual or 5000 cells/well for combinatorial drug studies were plated in a 96-well plate. Cells were then treated 24 hours later with temozolomide (Selleck Chem cat #S1237), olaparib (Selleck Chem #S1060), both drugs, or DMSO at an equivalent percent volume to the highest drug concentration. Cell viability was assayed 4 days later following a 12-hour incubation with alamarBlue (ThermoFisher cat #DAL1025) and detected using a fluorescence-based plate reader. For Zika virus studies, 5000 cells/well were infected with Dakar 41519 strain ZIKV at a multiplicity of infection of 5 (MOI 5) (74). Viability was assayed at 3 days of post infection using CellTiter-Glo according to manufacturer’s instructions.

## ACKNOWLEDGEMENTS

This research was supported by the National Cancer Institute funded Salk Institute Cancer Center (NIH/NCI CCSG: P30 014195) by a grant from Padres Pedal the Cause/RADY #PTC2019 (G.M), by a Pioneer Fund Postdoctoral Scholar Award (A.S), and the following NIH grants and fellowships: CA217066 (B.C.P.); CA217065 (R.C.G.); CA197718, CA238662, NS103434 (J.N.R.). G.M. was supported by the Frederick B. Rentschler Developmental Chair. Figures 1A and 7 were created with BioRender.com.

## SUPPLEMENTAL FIGURES

**Supplemental Figure S1:**
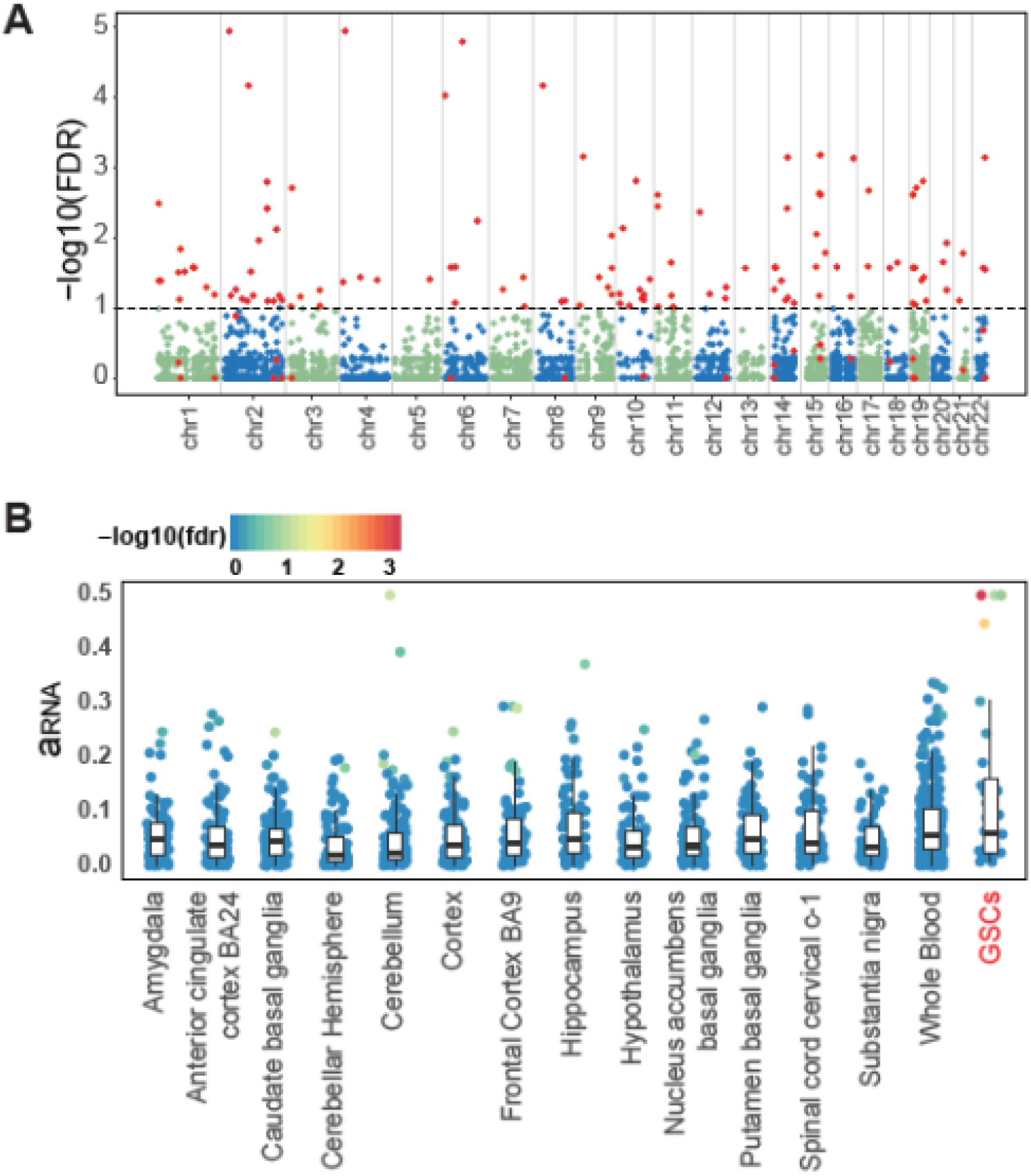
Recurrent allele-specific expression within glioblastoma stem cells. **A)** Manhattan plot demonstrating that genes with recurrent ASE in glioblastoma stem cells (GSCs) are not localized to any single genomic locus. The x-axis is the gene start position for all tested genes and y-axis is the −log10 transformed FDR corrected Fisher Exact Test p-value. Genes significant under an FDR of 10% are highlighted in red. **B)** Comparison of estimated ASE for *IP6K2* for GSCs, normal brain and whole-blood samples from GTEx.

**Supplemental Figure S2:**
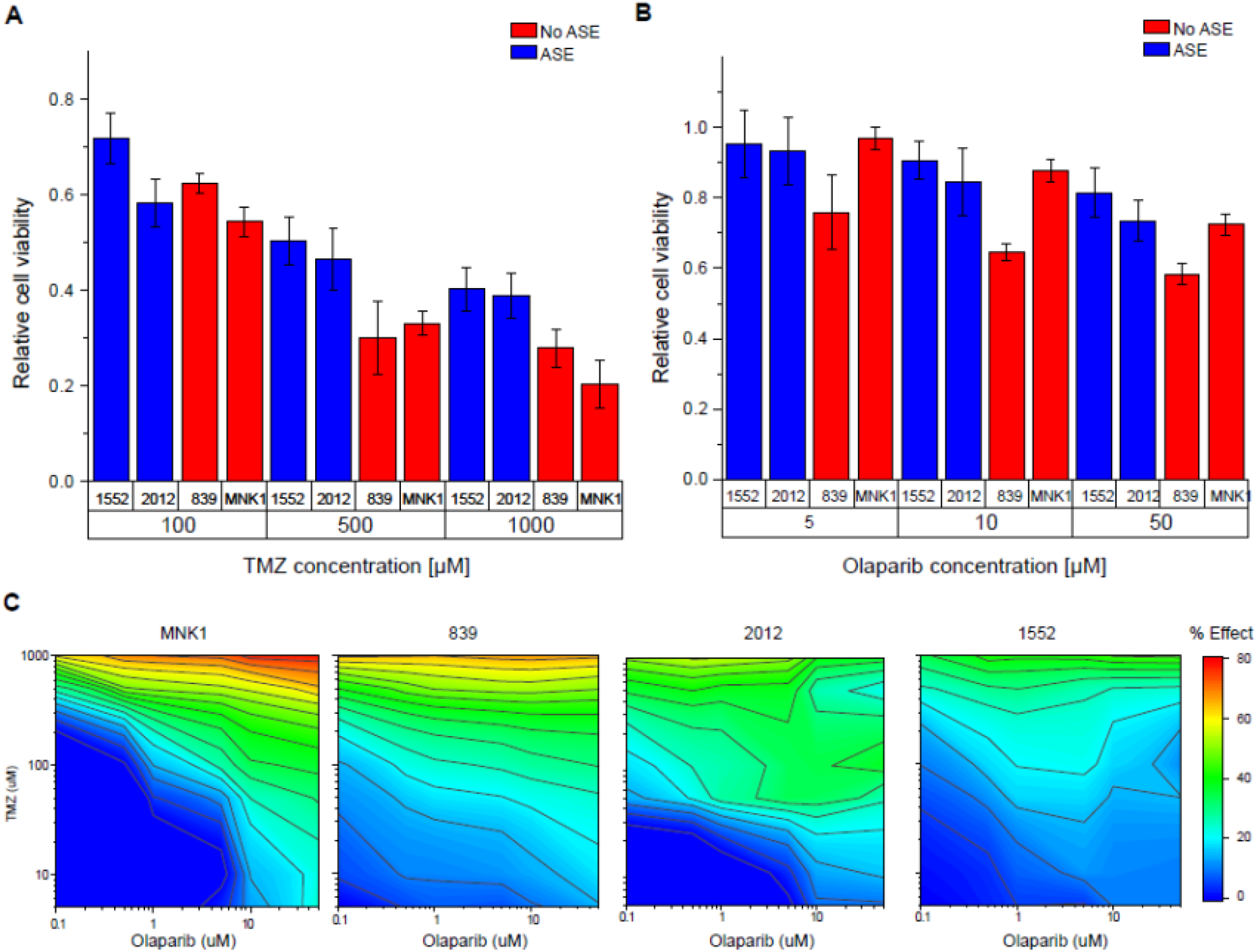
Cellular viability assays for GSCs. **A)** Relative cell viability plotted on the y-axis for each model treated with TMZ. Non-ASE models are in red and ASE models are in blue. **B)** Relative cell viability plotted on the y-axis for each model treated with olaparib. Non-ASE models are in red and ASE models are in blue. **C)** Cell viability relative to DMSO control is annotated on a rainbow scale with blue indicating high viability, or minimal drug effect, and red indicating low viability, or strong drug effect. Doses are scaled logarithmically on the x-axis (olaparib) and y axis (TMZ). Data are represented as mean +/− standard deviation. ASE: allele-specific expression; GSC: glioblastoma stem cell; KD: knockdown; OE: overexpression; TMZ: temozolomide.

**Supplemental Figure S3:**
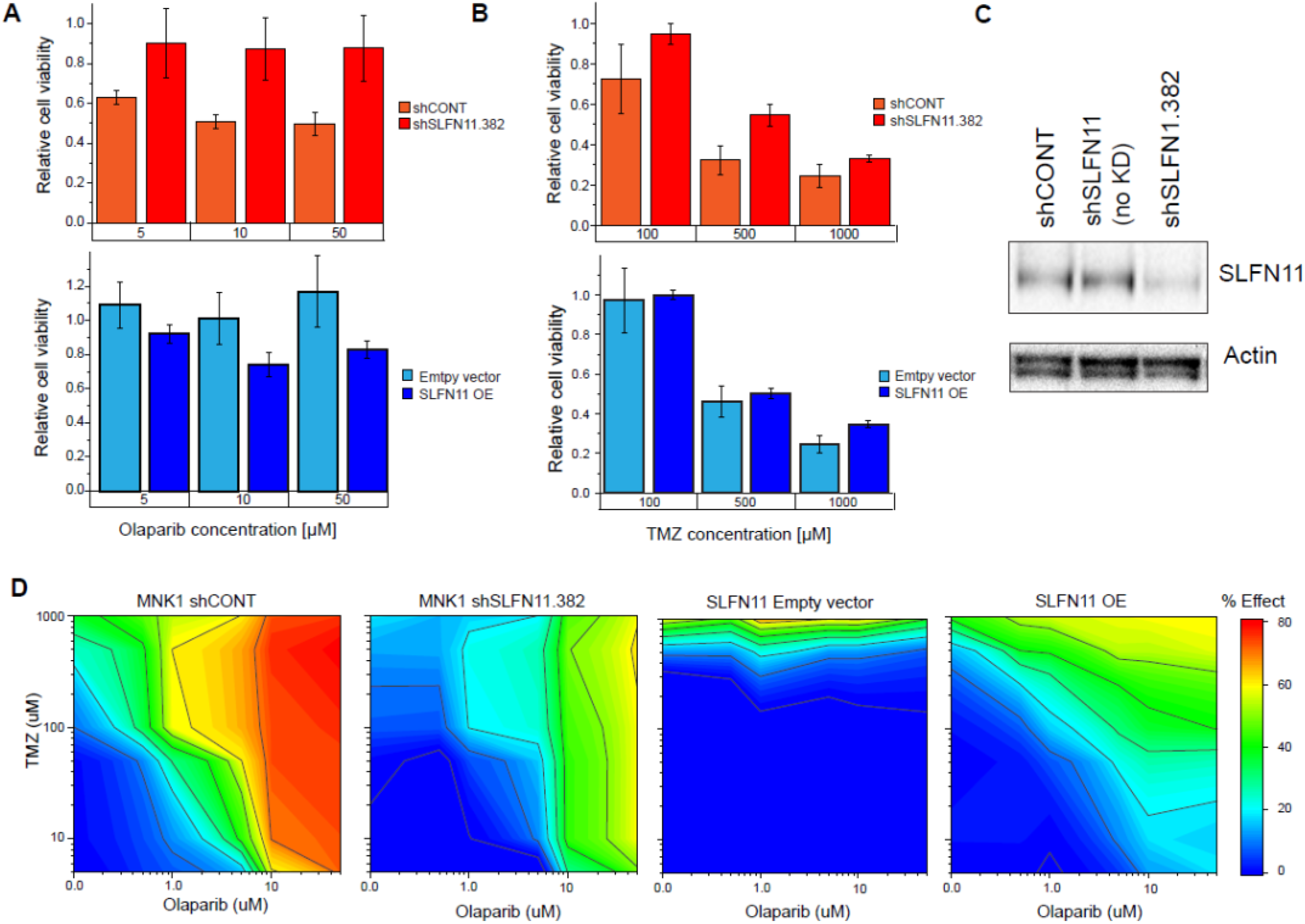
Cellular viability assays for GSCs with knockdown and overexpression of *SLFN11*. **A)** Relative cell viability plotted on the y-axis for each model treated with olaparib. Upper panel: MNK1 shCONT cells are in orange and SLFN11 KD in red; Lower panel: 2012 empty vector control cells are in light blue and SLFN11 OE are in blue. **B)** Relative cell viability plotted on the y-axis for each model treated with TMZ. Upper panel: MNK1 shCONT cells are in orange and SLFN11 KD in red; Lower panel: 2012 empty vector control cells are in light-blue and SLFN11 OE are in blue. **C)** Western blot of the MNK1 GSCs for SLFN11 and Actin expression. **D)** Cell viability relative to DMSO control is annotated on a rainbow scale with blue indicating high viability, or minimal drug effect, and red indicating low viability, or strong drug effect. Doses are scaled logarithmically on the x-axis (olaparib) and y axis (TMZ). Data are represented as mean +/− standard deviation. KD: knockdown; OE: overexpression; TMZ: temozolomide. Data are represented as mean +/− standard deviation. ASE: allele-specific expression; GSC: glioblastoma stem cell; KD: knockdown; OE: overexpression; TMZ: temozolomide.

**Supplemental Figure S4.**
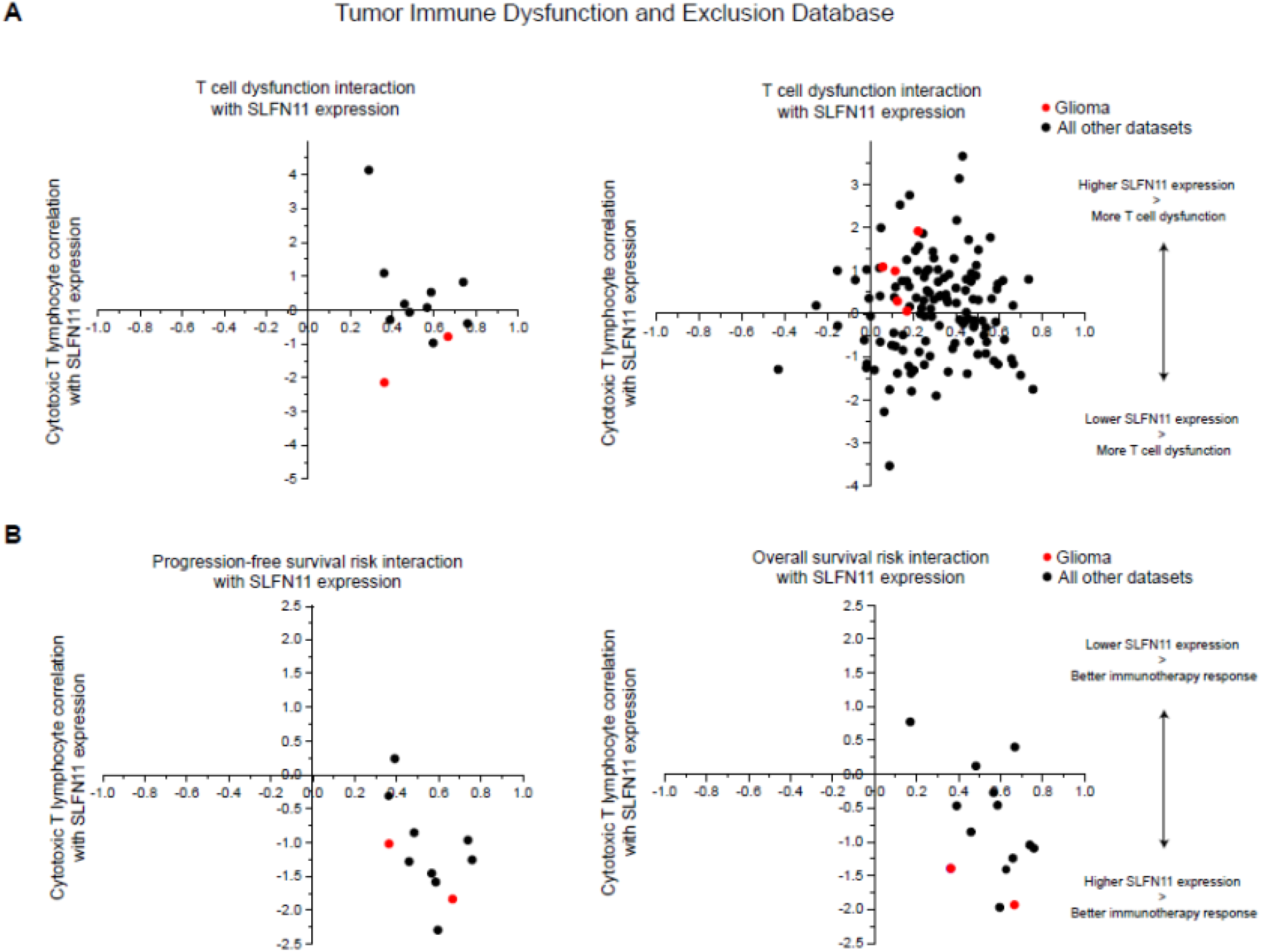
Dysfunction and Exclusion Database. **A)** Scatter plot showing positive interaction of *SLFN11* expression with T cell dysfunction (x-axis) and negative correlation of *SLFN11* expression with cytotoxic T lymphocytes (y-axis). Glioma samples are in red. **B)** Scatter plot demonstrating positive interaction of *SLFN11* expression with progression-free survival (x-axis) (left) or overall survival (right) vs negative correlation of *SLFN11* expression with cytotoxic T lymphocytes (y-axis). Glioma samples are in red.

## SUPPLEMENTAL TABLES

**Supplemental Table S1:** Results for ASE analysis for glioblastoma samples.

**Supplemental Table S2:** Results for Fisher Exact Test (FET) which was used for discovering genes which show enrichment of ASE in glioblastoma samples compared to whole blood from GTEx.

**Supplemental Table S3:** Results from Spearman’s correlation analysis between normalized gene expression and normalized coverage for H3K27ac ChIP-seq bins which are located within 3-100Kb of promoters of candidate genes.

**Supplemental Table S4:** Empirical p-value estimates for 29 *NOTCH1* CREs.

**Supplemental Table S5:** Results from Spearman’s correlation analysis between normalized gene expression and mean promoter methylation (β_promoter_).

## REFERENCES

1. Ostrom QT, Patil N, Cioffi G, Waite K, Kruchko C, Barnholtz-Sloan JS. CBTRUS Statistical Report: Primary Brain and Other Central Nervous System Tumors Diagnosed in the United States in 2013-2017. Neuro Oncol 2020;22:iv1–iv96

2. Snuderl M, Fazlollahi L, Le LP, Nitta M, Zhelyazkova BH, Davidson CJ, et al. Mosaic amplification of multiple receptor tyrosine kinase genes in glioblastoma. Cancer Cell 2011;20:810–7

3. Patel AP, Tirosh I, Trombetta JJ, Shalek AK, Gillespie SM, Wakimoto H, et al. Single-cell RNA-seq highlights intratumoral heterogeneity in primary glioblastoma. Science 2014;344:1396–401

4. Lathia JD, Mack SC, Mulkearns-Hubert EE, Valentim CL, Rich JN. Cancer stem cells in glioblastoma. Genes Dev 2015;29:1203–17

5. Chen J, Li Y, Yu TS, McKay RM, Burns DK, Kernie SG, et al. A restricted cell population propagates glioblastoma growth after chemotherapy. Nature 2012;488:522–6

6. Alvarado AG, Thiagarajan PS, Mulkearns-Hubert EE, Silver DJ, Hale JS, Alban TJ, et al. Glioblastoma Cancer Stem Cells Evade Innate Immune Suppression of Self-Renewal through Reduced TLR4 Expression. Cell Stem Cell 2017;20:450–61 e4

7. Eramo A, Ricci-Vitiani L, Zeuner A, Pallini R, Lotti F, Sette G, et al. Chemotherapy resistance of glioblastoma stem cells. Cell Death Differ 2006;13:1238–41

8. Bao S, Wu Q, McLendon RE, Hao Y, Shi Q, Hjelmeland AB, et al. Glioma stem cells promote radioresistance by preferential activation of the DNA damage response. Nature 2006;444:756–60

9. Brennan CW, Verhaak RG, McKenna A, Campos B, Noushmehr H, Salama SR, et al. The somatic genomic landscape of glioblastoma. Cell 2013;155:462–77

10. Nawaz Z, Patil V, Thinagararjan S, Rao SA, Hegde AS, Arivazhagan A, et al. Impact of somatic copy number alterations on the glioblastoma miRNome: miR-4484 is a genomically deleted tumour suppressor. Mol Oncol 2017;11:927–44

11. Munoz-Hidalgo L, San-Miguel T, Megias J, Monleon D, Navarro L, Roldan P, et al. Somatic copy number alterations are associated with EGFR amplification and shortened survival in patients with primary glioblastoma. Neoplasia 2020;22:10–21

12. Cao S, Zhou DC, Oh C, Jayasinghe RG, Zhao Y, Yoon CJ, et al. Discovery of driver non-coding splice-site-creating mutations in cancer. Nat Commun 2020;11:5573

13. Puente XS, Bea S, Valdes-Mas R, Villamor N, Gutierrez-Abril J, Martin-Subero JI, et al. Non-coding recurrent mutations in chronic lymphocytic leukaemia. Nature 2015;526:519–24

14. Flavahan WA, Drier Y, Liau BB, Gillespie SM, Venteicher AS, Stemmer-Rachamimov AO, et al. Insulator dysfunction and oncogene activation in IDH mutant gliomas. Nature 2016;529:110–4

15. Liu Y, Li C, Shen S, Chen X, Szlachta K, Edmonson MN, et al. Discovery of regulatory noncoding variants in individual cancer genomes by using cis-X. Nat Genet 2020;52:811–8

16. Huang FW, Hodis E, Xu MJ, Kryukov GV, Chin L, Garraway LA. Highly recurrent TERT promoter mutations in human melanoma. Science 2013;339:957–9

17. Horn S, Figl A, Rachakonda PS, Fischer C, Sucker A, Gast A, et al. TERT promoter mutations in familial and sporadic melanoma. Science 2013;339:959–61

18. Mansour MR, Abraham BJ, Anders L, Berezovskaya A, Gutierrez A, Durbin AD, et al. Oncogene regulation. An oncogenic super-enhancer formed through somatic mutation of a noncoding intergenic element. Science 2014;346:1373–7

19. Gasperini M, Hill AJ, McFaline-Figueroa JL, Martin B, Kim S, Zhang MD, et al. A Genome-wide Framework for Mapping Gene Regulation via Cellular Genetic Screens. Cell 2019;176:377–90 e19

20. Pirinen M, Lappalainen T, Zaitlen NA, Consortium GT, Dermitzakis ET, Donnelly P, et al. Assessing allele-specific expression across multiple tissues from RNA-seq read data. Bioinformatics 2015;31:2497–504

21. Mohammadi P, Castel SE, Cummings BB, Einson J, Sousa C, Hoffman P, et al. Genetic regulatory variation in populations informs transcriptome analysis in rare disease. Science 2019;366:351–6

22. Pastinen T. Genome-wide allele-specific analysis: insights into regulatory variation. Nat Rev Genet 2010;11:533–8

23. Skelly DA, Johansson M, Madeoy J, Wakefield J, Akey JM. A powerful and flexible statistical framework for testing hypotheses of allele-specific gene expression from RNA-seq data. Genome Res 2011;21:1728–37

24. Buckberry S, Bianco-Miotto T, Hiendleder S, Roberts CT. Quantitative allele-specific expression and DNA methylation analysis of H19, IGF2 and IGF2R in the human placenta across gestation reveals H19 imprinting plasticity. PLoS One 2012;7:e51210

25. Deng Q, Ramskold D, Reinius B, Sandberg R. Single-cell RNA-seq reveals dynamic, random monoallelic gene expression in mammalian cells. Science 2014;343:193–6

26. Mack SC, Singh I, Wang X, Hirsch R, Wu Q, Villagomez R, et al. Chromatin landscapes reveal developmentally encoded transcriptional states that define human glioblastoma. J Exp Med 2019;216:1071–90

27. Rachmilewitz J, Goshen R, Ariel I, Schneider T, de Groot N, Hochberg A. Parental imprinting of the human H19 gene. FEBS Lett 1992;309:25–8

28. Ma Y, Gong Y, Cheng Z, Loganathan S, Kao C, Sarkaria JN, et al. Critical functions of RhoB in support of glioblastoma tumorigenesis. Neuro Oncol 2015;17:516–25

29. Baldwin RM, Parolin DA, Lorimer IA. Regulation of glioblastoma cell invasion by PKC iota and RhoB. Oncogene 2008;27:3587–95

30. Morrison BH, Bauer JA, Kalvakolanu DV, Lindner DJ. Inositol hexakisphosphate kinase 2 mediates growth suppressive and apoptotic effects of interferon-beta in ovarian carcinoma cells. J Biol Chem 2001;276:24965–70

31. Chakraborty A, Koldobskiy MA, Sixt KM, Juluri KR, Mustafa AK, Snowman AM, et al. HSP90 regulates cell survival via inositol hexakisphosphate kinase-2. Proc Natl Acad Sci U S A 2008;105:1134–9

32. Abraham BJ, Hnisz D, Weintraub AS, Kwiatkowski N, Li CH, Li Z, et al. Small genomic insertions form enhancers that misregulate oncogenes. Nat Commun 2017;8:14385

33. Corces MR, Granja JM, Shams S, Louie BH, Seoane JA, Zhou W, et al. The chromatin accessibility landscape of primary human cancers. Science 2018;362

34. Hegi ME, Diserens AC, Gorlia T, Hamou MF, de Tribolet N, Weller M, et al. MGMT gene silencing and benefit from temozolomide in glioblastoma. N Engl J Med 2005;352:997–1003

35. Zoppoli G, Regairaz M, Leo E, Reinhold WC, Varma S, Ballestrero A, et al. Putative DNA/RNA helicase Schlafen-11 (SLFN11) sensitizes cancer cells to DNA-damaging agents. Proc Natl Acad Sci U S A 2012;109:15030–5

36. Mu Y, Lou J, Srivastava M, Zhao B, Feng XH, Liu T, et al. SLFN11 inhibits checkpoint maintenance and homologous recombination repair. EMBO Rep 2016;17:94–109

37. Lok BH, Gardner EE, Schneeberger VE, Ni A, Desmeules P, Rekhtman N, et al. PARP Inhibitor Activity Correlates with SLFN11 Expression and Demonstrates Synergy with Temozolomide in Small Cell Lung Cancer. Clin Cancer Res 2017;23:523–35

38. Ianevski A, Giri AK, Aittokallio T. SynergyFinder 2.0: visual analytics of multi-drug combination synergies. Nucleic Acids Res 2020;48:W488–W93

39. Li M, Kao E, Gao X, Sandig H, Limmer K, Pavon-Eternod M, et al. Codon-usage-based inhibition of HIV protein synthesis by human schlafen 11. Nature 2012;491:125–8

40. Malone D, Lardelli RM, Li M, David M. Dephosphorylation activates the interferon-stimulated Schlafen family member 11 in the DNA damage response. J Biol Chem 2019;294:14674–85

41. Valdez F, Salvador J, Palermo PM, Mohl JE, Hanley KA, Watts D, et al. Schlafen 11 Restricts Flavivirus Replication. J Virol 2019;93

42. Alonso MM, Jiang H, Gomez-Manzano C, Fueyo J. Targeting brain tumor stem cells with oncolytic adenoviruses. Methods Mol Biol 2012;797:111–25

43. Martuza RL, Malick A, Markert JM, Ruffner KL, Coen DM. Experimental therapy of human glioma by means of a genetically engineered virus mutant. Science 1991;252:854–6

44. Zhu Z, Gorman MJ, McKenzie LD, Chai JN, Hubert CG, Prager BC, et al. Zika virus has oncolytic activity against glioblastoma stem cells. J Exp Med 2017;214:2843–57

45. Wang Q, Hu B, Hu X, Kim H, Squatrito M, Scarpace L, et al. Tumor Evolution of Glioma-Intrinsic Gene Expression Subtypes Associates with Immunological Changes in the Microenvironment. Cancer Cell 2017;32:42–56 e6

46. Liberzon A, Birger C, Thorvaldsdottir H, Ghandi M, Mesirov JP, Tamayo P. The Molecular Signatures Database (MSigDB) hallmark gene set collection. Cell Syst 2015;1:417–25

47. Yi M, Nissley DV, McCormick F, Stephens RM. ssGSEA score-based Ras dependency indexes derived from gene expression data reveal potential Ras addiction mechanisms with possible clinical implications. Sci Rep 2020;10:10258

48. Subramanian A, Tamayo P, Mootha VK, Mukherjee S, Ebert BL, Gillette MA, et al. Gene set enrichment analysis: a knowledge-based approach for interpreting genome-wide expression profiles. Proc Natl Acad Sci U S A 2005;102:15545–50

49. Silginer M, Nagy S, Happold C, Schneider H, Weller M, Roth P. Autocrine activation of the IFN signaling pathway may promote immune escape in glioblastoma. Neuro Oncol 2017;19:1338–49

50. Delbare SYN, Clark AG. Allele-specific expression elucidates cis-regulatory logic. PLoS Genet 2018;14:e1007690

51. Mayba O, Gilbert HN, Liu J, Haverty PM, Jhunjhunwala S, Jiang Z, et al. MBASED: allele-specific expression detection in cancer tissues and cell lines. Genome Biol 2014;15:405

52. Shames DS, Minna JD. IP6K2 is a client for HSP90 and a target for cancer therapeutics development. Proc Natl Acad Sci U S A 2008;105:1389–90

53. Koldobskiy MA, Chakraborty A, Werner JK, Jr., Snowman AM, Juluri KR, Vandiver MS, et al. p53-mediated apoptosis requires inositol hexakisphosphate kinase-2. Proc Natl Acad Sci U S A 2010;107:20947–51

54. Borghese L, Dolezalova D, Opitz T, Haupt S, Leinhaas A, Steinfarz B, et al. Inhibition of notch signaling in human embryonic stem cell-derived neural stem cells delays G1/S phase transition and accelerates neuronal differentiation in vitro and in vivo. Stem Cells 2010;28:955–64

55. Ables JL, Decarolis NA, Johnson MA, Rivera PD, Gao Z, Cooper DC, et al. Notch1 is required for maintenance of the reservoir of adult hippocampal stem cells. J Neurosci 2010;30:10484–92

56. Li J, Cui Y, Gao G, Zhao Z, Zhang H, Wang X. Notch1 is an independent prognostic factor for patients with glioma. J Surg Oncol 2011;103:813–7

57. Purow BW, Haque RM, Noel MW, Su Q, Burdick MJ, Lee J, et al. Expression of Notch-1 and its ligands, Delta-like-1 and Jagged-1, is critical for glioma cell survival and proliferation. Cancer Res 2005;65:2353–63

58. Hu YY, Zheng MH, Cheng G, Li L, Liang L, Gao F, et al. Notch signaling contributes to the maintenance of both normal neural stem cells and patient-derived glioma stem cells. BMC Cancer 2011;11:82

59. Zhang XP, Zheng G, Zou L, Liu HL, Hou LH, Zhou P, et al. Notch activation promotes cell proliferation and the formation of neural stem cell-like colonies in human glioma cells. Mol Cell Biochem 2008;307:101–8

60. Man J, Yu X, Huang H, Zhou W, Xiang C, Huang H, et al. Hypoxic Induction of Vasorin Regulates Notch1 Turnover to Maintain Glioma Stem-like Cells. Cell Stem Cell 2018;22:104–18 e6

61. Hai L, Zhang C, Li T, Zhou X, Liu B, Li S, et al. Notch1 is a prognostic factor that is distinctly activated in the classical and proneural subtype of glioblastoma and that promotes glioma cell survival via the NF-kappaB(p65) pathway. Cell Death Dis 2018;9:158

62. Fassl A, Tagscherer KE, Richter J, Berriel Diaz M, Alcantara Llaguno SR, Campos B, et al. Notch1 signaling promotes survival of glioblastoma cells via EGFR-mediated induction of anti-apoptotic Mcl-1. Oncogene 2012;31:4698–708

63. Luan J, Gao X, Hu F, Zhang Y, Gou X. SLFN11 is a general target for enhancing the sensitivity of cancer to chemotherapy (DNA-damaging agents). J Drug Target 2020;28:33–40

64. Fernandes C, Costa A, Osorio L, Lago RC, Linhares P, Carvalho B, et al. Current Standards of Care in Glioblastoma Therapy. In: De Vleeschouwer S, editor. Glioblastoma. Brisbane (AU)2017.

65. Houtgast EJ, Sima VM, Bertels K, Al-Ars Z. Hardware acceleration of BWA-MEM genomic short read mapping for longer read lengths. Comput Biol Chem 2018;75:54–64

66. Li H, Handsaker B, Wysoker A, Fennell T, Ruan J, Homer N, et al. The Sequence Alignment/Map format and SAMtools. Bioinformatics 2009;25:2078–9

67. Dobin A, Davis CA, Schlesinger F, Drenkow J, Zaleski C, Jha S, et al. STAR: ultrafast universal RNA-seq aligner. Bioinformatics 2013;29:15–21

68. Liao Y, Smyth GK, Shi W. featureCounts: an efficient general purpose program for assigning sequence reads to genomic features. Bioinformatics 2014;30:923–30

69. Love MI, Huber W, Anders S. Moderated estimation of fold change and dispersion for RNA-seq data with DESeq2. Genome Biol 2014;15:550

70. van de Geijn B, McVicker G, Gilad Y, Pritchard JK. WASP: allele-specific software for robust molecular quantitative trait locus discovery. Nat Methods 2015;12:1061–3

71. Alexa A, Rahnenfuhrer J, Lengauer T. Improved scoring of functional groups from gene expression data by decorrelating GO graph structure. Bioinformatics 2006;22:1600–7

72. Zhang Y, Liu T, Meyer CA, Eeckhoute J, Johnson DS, Bernstein BE, et al. Model-based analysis of ChIP-Seq (MACS). Genome Biol 2008;9:R137

73. Yu G, Wang LG, He QY. ChIPseeker: an R/Bioconductor package for ChIP peak annotation, comparison and visualization. Bioinformatics 2015;31:2382–3

74. Sapparapu G, Fernandez E, Kose N, Bin C, Fox JM, Bombardi RG, et al. Neutralizing human antibodies prevent Zika virus replication and fetal disease in mice. Nature 2016;540:443–7

